# Metal-Assisted and Microwave-Accelerated Germination

**DOI:** 10.1101/743252

**Authors:** Kadir Aslan, Monet Stevenson, Janelle Guy, Enock Bonyi, Muzaffer Mohammed, Birol Ozturk, Kyle Drake, Freeman McLean, Ashley Souffrant, Amber Bigio

**Author notes:** Corresponding Author, Voice.

## Abstract

We report the proof-of-principle demonstration of a methodology, called Metal-Assisted and Microwave-Accelerated Germination, to modulate the germination of plant seeds and growth of plants using gold nanoparticles (Au NPs) and microwave heating. As a model plant seed, basil seeds were heated in a solution of 20 nm Au NPs using a microwave waveguide fiber connected to a solid-state microwave operating at 8 GHz at 20 W, which resulted in the development of longer basil gum as observed by optical microscopy. In control experiments, Au NPs or microwave heating was omitted to establish a baseline growth level under standard experimental conditions (no microwave heating or no Au NPs). Our results also show that hydroponic growth and soil growth of basil plants can be delayed with the use of 20 nm Au NPs at room temperature without microwave heating. The combined use of 20 nm Au NPs and microwave heating at 10 W for 6 minutes results in accelerated growth prolonged life of basil plants.

## INTRODUCTION

As humanity continue to face food shortage issues,(Perfecto and Vandermeer, 2010; Clay, 2011; Crist *et al.*, 2017) a number of methods are developed with the aim of greater food production(Townsend and Porder, 2012; Scott *et al.*, 2018; Jacobsen *et al.*, 2019; Tiberius *et al.*, 2019). To increase the production of food from plants, their seeds are subjected to a number of physical methods, such as, ultraviolet light treatment,(Noble, 2002) magnetic field treatment,(Maffei, 2014) and hot water soaking,(Hsu *et al.*, 2003) and chemical methods(Farajollahi *et al.*, 2014) with the specific goal of reducing the natural germination times of seeds. The above-mentioned methods are either time-consuming, labor-intensive, or produce harmful by-product; thus, alternative techniques for the acceleration of the plant germination process are being pursued. Recently, emerging techniques, such as, cold plasma treatment,(Ling *et al.*, 2014; de Groot *et al.*, 2018) radio frequency,(Mildažienė *et al.*, 2019) or a combined use of the three(Mildaziene *et al.*, 2016) were developed for the potential rapid germination of plant seeds. Although these techniques enhance the germination process, several limitations including lack of reduction of germination time, inconsistent results, and unsuitable moisture around the seeds hinder their wide-spread application.

In addition, carbon-based and metallic nanoparticles (NPs) have been shown to enhance germination and seedling growth, physiological activities, nitrogen metabolism, mRNA expression and protein expression levels, and a few examples are summarized here. For example, in a study by Kole *et. al.* (Kole *et al.*, 2013) fullerol was used to treat seeds to evaluate its effect on water uptake and fruit biomass, where an increase of up to 54% in biomass yield and 24% in water content was reported. Mahakham *et. al.* (Mahakham *et al.*, 2017) primed rice seeds with phytosynthesized silver (Ag) NPs improved germination and seedling vigor when compared to an unprimed control, silver nitrate priming, and conventional hydropriming. A key observation of this study is the production of more reactive oxygen species in germinating seeds undergoing nano-priming treatment. Mahakham *et. al.* (Mahakham *et al.*, 2017) proposed several mechanisms for germinations process, which includes creation of nanopores for water uptake, rebooting of ROS antioxidant systems in seeds, generation of hydroxyl radicals for the loosening of cell wall, and formation of nanocatalyst for starch hydrolysis fastening. Thuesombat *et. al.* (Thuesombat *et al.*, 2014) investigated the possible effects of different sized Ag NPs (20 nm, 30-60 nm, 70-120 nm, and 150 nm in diameter) on jasmine rice (*Oryza sativa*) seed germination and seedling growth, where Ag NPs > 150 nm in size were determined to have a more negative effect on rice seedling growth than smaller sizes. Greater permeation of Ag NPs is found in roots when smaller Ag NPs (<150 nm) are used, yet the smaller Ag NPs were transported to the shoots with less effectively than larger sizes. The use of Ag NPs yields inconsistent and negative results on the growth and development of plants and has the potential to increase environmental toxicity. The shortcomings of existing methods limit ability to accurately and reliably test with sensitivity, specificity, and rapidity. Efforts to address these inadequacies will require innovative techniques capable of rapid germination of seeds.

Our research group has been investigating the effect of combined use of metal NPs and microwave heating on crystallization of amino acids (Pinard and Aslan, 2010) and proteins,(Mauge-Lewis *et al.*, 2015) decrystallization of uric acid crystals,(Thompson *et al.*, 2017; Boone-Kukoyi *et al.*, 2019) and biosensors,(Abel *et al.*, 2014; Mohammed *et al.*, 2014). In these studies, metal NPs are either immobilized on a planar surface to assist in the attachment of biological materials through chemisorption (for crystallization (Pinard and Aslan, 2010) and biosensors (Abel *et al.*, 2014; Mohammed *et al.*, 2014)) or in solution to function as “nano-bullets” in decrystallization of crystals on a planar surface (Thompson *et al.*, 2017; Boone-Kukoyi *et al.*, 2019). Based on our understanding of combined use of metal NPs and microwave heating to accelerate physical and biological interactions, we hypothesized that the germination of plant seeds and subsequent growth of plants can be modulated by the use of Au NPs and microwave heating.

In this proof-of-principle work, we report a methodology to modulate (i.e., delay or accelerate) the germination of basil seeds and subsequent growth of basil plant based on combined use of Au NPs and microwave heating, called Metal-Assisted and Microwave-Accelerated Germination (MAMAG). The crux of the MAMAG technique is depicted in **Figure 1**. Citrate stabilized (negatively-charged) Au NPs in solution are accelerated by the incident microwave field towards the basil seeds. The increased kinetic energy of the Au NPs increases collisions and interactions between NPs and the basil seeds. Subsequently, the following series of events are thought to occur: the seed testae develop cracks and ridges due to collisions with Au NPs and microwave heating of water contained in the basil seeds (i.e., due to resistive losses), Au NPs bind to enzymes on the seed surface, enzyme activity on the basil seeds are modified by Au NPs and the germination process can be delayed or accelerated based on the extent of enzyme activity that is directly correlated to the presence of Au NPs and microwave heating. We report that the germination of basil seeds, hydroponic growth and soil growth of basil seeds can be delayed with the use of 20 nm Au NPs without microwave heating and accelerated in combination with microwave heating at up to 20 W with a solid-state microwave source operating at 8 GHz. Since previous reports showed that larger Ag NPs had a negative effect on the growth of plants, we did not attempt to repeat these experiments with Au NPs in our study.

**Figure 1.**
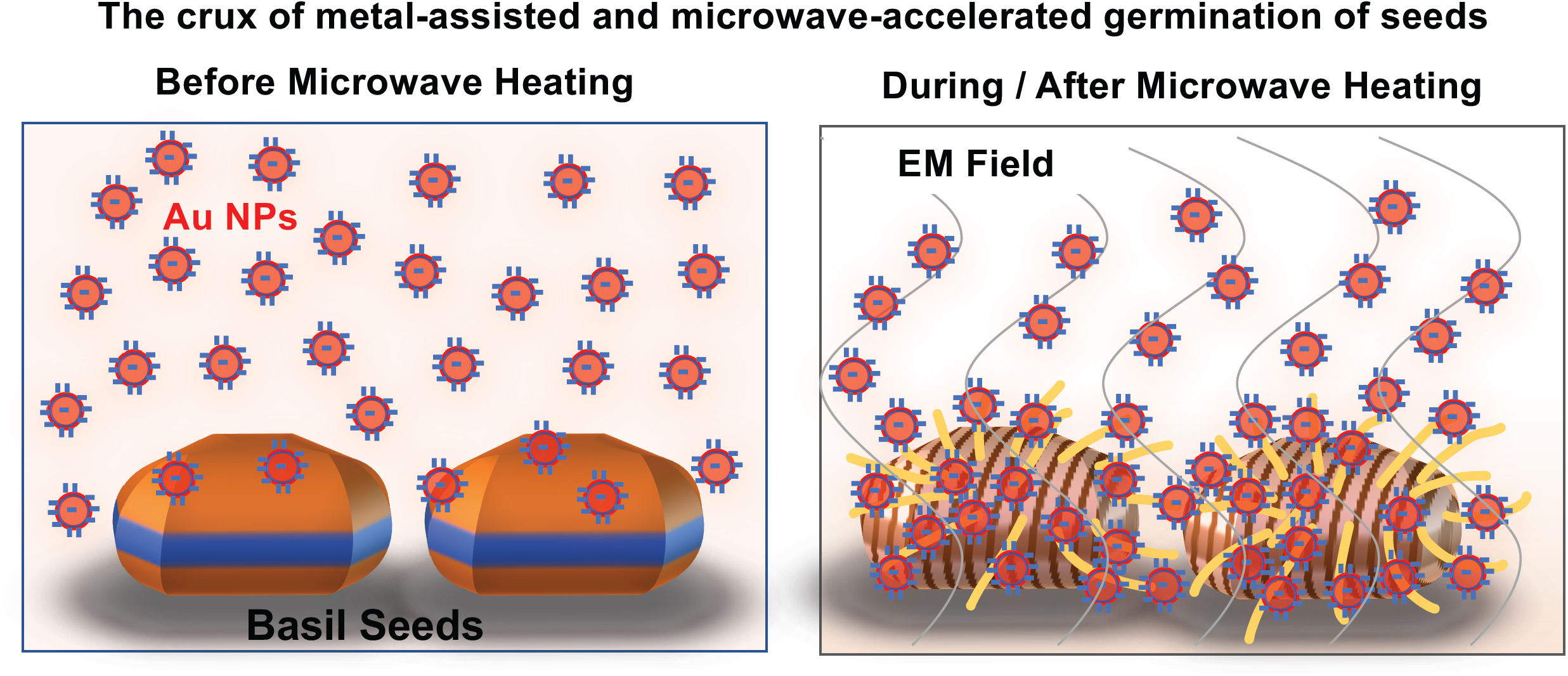
The crux of the MAMAG technique: citrate-stabilized (negatively charged) gold nanoparticles (Au NPs) are accelerated by the electromagnetic field. The increased kinetic energy of Au NPs increases collisions between nanoparticles and basil seeds. Seed testae develop cracks and ridges due to collisions of Au NPs with the seed surface and resistive losses due to microwave heating of water within the seeds, leading to controlled germination of seeds. Esterase activity on the seed surface is also increased due to the presence of Au NPs and microwave heating. Gold nanoparticles bind to the enzymes/proteins via Au-thiol and Au/primary amine bonds and electrostatic interactions. Wavelength of monomode electromagnetic field at 8 GHz is ca. 3.75 cm.

## MATERIALS and METHODS

### Materials

*Ocimum basilicum* (basil) seeds were purchased from a local seed store in Baltimore, Maryland, USA in 2014 and were kept in cool and dry place until use (i.e., basil seeds were 4 years old at the time of their use in 2018). Citrate-stabilized 20 nm Au NPs were purchased from Sigma Aldrich (Milwaukee, WI, USA) and stored at 4°C. 4-Methylumberlliferyl butyrate (product number: 19362-5G) was purchased from Sigma Aldrich (USA) and was used as fluorogenic substrates for esterase. HTS well plates containing 96, flat bottomed wells, was acquired from Sarstedt (Newton, NC, USA, product number: 82.1583.001).

### Instrumentation

A Digi-microscope from Digitech Industries, Inc. (USA) was used to view basil seeds during microwave heating using the ISYS800 system (an 8 GHz solid-state microwave source with variable power levels from 0-20 W, from Emblation Ltd (Scotland, UK). Microwave heating of samples is accomplished using a ceramic applicator tip (diameter = 5 mm) and a coaxial cable (outer diameter = 6.7 mm and length = 1.3 m) that delivers microwaves from the source to the applicator tip. The natural luminescence of the seeds and the esterase activity was measured using a Cytation™ 5 Cell Imaging Multi-Mode Reader from BioTek Instruments, Inc. (Winooski, VT, USA). Changes in temperature of the seeds during microwave heating were measured using the ETS320 model thermal imaging camera from FLIR^®^Systems, Inc. (Wilsonville, OR, USA), which was placed directly under the optically clear HTS wells. A Phenom XL PhenomWorld Desktop Scanning Electron Microscope (SEM) equipped with elemental analysis capability (ThermoFisher Scientific, Waltham, MA, USA) was used to image the seeds before and after microwave heating.

Micro-seed hydroponic growth pads and pH Perfect Technology Grow nutrient prepared by Advanced Nutrients (Abbotsford, BC, Canada, optimum dilution 1:1000) were utilized in hydroponic studies. The soil used to fill growth pads was obtained Miracle Gro Seed Starting Potting Mix (EcoGreenText, Inc. NY, USA). The greenhouse (dimensions 120 × 120 × 200 cm) was purchased from (Amazon.com) and a light source capable of reflecting all light was obtained from Amazon.com. All water used was purified using a Millipore Direct Q3 apparatus.

## Methods

### Determination of seed population viability at room temperature

One hundred basil seeds were completely covered with water in an open top pan at room temperature for 1 hour. Then, basil seeds were placed on top of 0.22 μm filter paper in a standard circular petri dish. Petri dish was closed with its original lid and were kept in the dark for 4 days during which the germination percentage was calculated daily. The percentage of basil seed germination was calculated using the following equation: ([# of seeds germinated/total #of seeds] × 100) to estimate the viability of the seed population.

### Determination of the combined effect of gold nanoparticles and microwave heating on basil seeds

Basil seeds were placed in either 50 μL deionized water or 50 μL 20 nm gold nanoparticles (Au NPs) in a 96-well flat bottom plate. Each well of the HTS plates can hold up to 100 μL of solvent. Head space in the HTS wells after the addition of solutions to basil seeds was used to allow to help cope with the increase in water vapor pressure and to prevent spillage of water or 20 nm Au NPs from the wells during microwave heating. Dry basil seeds without solution were used as control seeds. All basil seeds (i.e., dry basil seeds, basil seeds in water, and basil seeds in 20 nm Au NPs) were treated with continuous microwave heating (2 W, 10 W, 20 W) for 6 minutes. Before and after microwave heating, brightfield, fluorescent images, and SEM images were obtained to characterize the effect of microwave heating on basil seeds. Fluorescent images were obtained using a DAPI (blue) filter, a GFP (green) filter, and an RFP (red) filter setting of our Cytation 5 Multi-mode Cell Imager. Basil seed gum (mucilage) was characterized by SEM after microwave heating. We also measured temperature changes in the HTS wells with basil seeds during microwave heating using thermal imaging. Control experiments without microwave heating were performed at room temperature.

### Determination of gold nanoparticles and microwave heating effects on hydroponic growth

We utilized a modified flood and drain hydroponic growth system to determine the combined effect of microwave heating and 20 nm Au NPs on hydroponic growth. In our hydroponic growth system, a polypropylene hydroponic growth pad was placed inside a petri dish. Three basil seeds kept in water at room temperature for 6 minutes (control experiment, no microwave heating), basil seeds kept in a solution of 20 nm Au NPs at room temperature for 6 minutes (no microwave heating), basil seeds in water with continuous microwave heating at 10 W for 6 minutes, and basil seeds in a solution of 20 nm Au NPs with continuous microwave heating at 10 W for 6 minutes were placed in the hydroponic growth pad. Holes in the hydroponic growth pad were made as same depth as the basil seeds to prevent root entanglement. The hydroponic growth system was well hydrated without evaporation of water throughout basil seed germination process.

### Determination of gold nanoparticles and microwave heating effects on growth of basil seeds in soil in a greenhouse

Basil seeds were placed in either 50 μL deionized water or 50 μL solution of 20 nm Au NPs in a 96-well flat bottom plate. All basil seeds (i.e., basil seeds in water and basil seeds in 20 nm Au NPs) were treated with continuous microwave heating (2 W, 10 W, 20 W) for 6 minutes. After microwave heating, basil seeds were placed in seedling starter trays. Seedling starting trays with basil seeds were placed in a greenhouse with LED light to monitor growth. Basil seeds kept at room temperature and in water at 35°C without microwave heating for 6 minutes were used as control seeds. All basil seeds were exposed to LED lighting system (Bloomspect, China, model BS300, 300 W) in 8-hour on and off cycles.

### Thermal Imaging Analysis

A FLIR thermal imaging camera (model ETS320) was used to determine the changes in temperature in the solution containing basil seed during microwave heating and at room temperature (control experiment, no microwave heating). Basil seeds were exposed to the treatment of no solution (dry), water, or 20 nm Au NPs with no microwave treatment or 2 W, 10 W, or 20 W of microwave treatment for 6 minutes while simultaneously recording with the thermal imaging camera. Thermal images were taken every 30 seconds using FLIR^®^ Systems software.

### Scanning Electron Microscopy (SEM)

The topography of basil seeds prior to and after exposure to experimental conditions described in the previous section was studied with SEM. Untreated basil seeds were placed on a piece of carbon tape and loaded into a Phenom XL SEM instrument. After focusing and setting the magnification to 300x, an image of the seed was captured. The same basil seeds were then subjected to the treatment of no solution (dry), water, or 20 nm Au NPs with no microwave heating or 2 W, 10 W, or 20 W of microwave heating for 6 minutes. The seed was then reloaded into the SEM machine and another image was captured. Elemental analysis on the captured images were carried out to quantify the extent of 20 nm Au NPs on the basil seeds.

### Measurement of Electrical Conductivity of Basil Seeds

Keithley source meter (model 2450) was used to measure electrical conductivity of basil seeds soaked either in water or in a solution of 20 nm Au NPs. Dry basil seeds were used as control experiment. Using the graphic user interphase window of the instrument, basic source measure settings were set as follows; measurement type = ohmmeter, sense = 2-wire sense, range of resistance = auto, and current source parameters (range =auto, source = 1 μA and voltage = 21 V). The two wire terminals from the source meter were pressed on each side of the seeds and the resistance read-out was recorded. Assuming that the seeds were elliptical in shape, the longer (1.1 mm) and shorter (0.11 mm) lengths were measured, and the cross-sectional area determined. Resistivity values were calculated using resistance, area and length measurements. Electrical conductivity was calculated as a reciprocal of resistivity.

### Finite-Difference Time-Domain (FDTD) Simulations

FDTD electromagnetic simulations were performed to determine the percentage of microwave absorption by each component of the system and to visualize the electric field propagation through the structure. MIT’s open source MEEP FDTD software (Oskooi *et al.*, 2010) was utilized for the two-dimensional simulations. Dielectric constants of seeds at 8 GHz microwave frequency were used in all simulations.(Gabriel *et al.*, 1996) Basil seeds were modeled as an elliptical object (1.1 mm in length and 0.11 in height) placed in air or water. In the electric field visualization simulations, a monomode 8 GHz microwave radiation was modeled as a fixed frequency continuous source located on the top part of the simulation cell. As in the experiments, the microwave radiation was transmitted to the structure through a 5 mm diameter waveguide, enabling single mode transmission. The field images depicted the propagation of the microwave radiation through the structure.

Resistive losses within the basil seeds caused by the applied electromagnetic field were predicted using COMSOL software (Boston, MA, USA). In this regard, basil seeds were modeled as a 1.1 mm diameter round object placed in water (fully immersed) and were placed inside a structure with the size of a high-throughput screening well. A monomode microwave source operating at 8 GHz was placed on top of the well structure to completely cover the well similar to the experimental setup. Resistive losses (in W/m^3^) within the basil seeds were calculated and is shown in three or twelve slices starting from the bottom of the seeds to the top of the seeds. The number of basil seeds used in FDTD simulations (1, 3, 4, and 10 basil seeds) were used to demonstrate the homogeneous heating of all seeds and were similar to that was obtained for a single basil seed.

## RESULTS AND DISCUSSION

### Determination of Seed Population Viability

To assess and minimize the potential experimental errors in germination due to inherent ability of basil seeds purchased from the vendor and the 4 years of storage, overall viability of basil seeds at room temperature was determined prior to the commencement of all experiments. In this regard, the percentage of viable basil seeds in 100 basil seeds was determined for up to 4 days (4-7 days are required for germination of basil seeds at room temperature). Fifteen percent (n = 15) of the basil seeds germinated at day 2. At day 3, 85% (n = 85) of the basil seeds sprouted. Nighty-eight percent (n = 98) of the basil seeds germinated within 4 days. and only 2% of the basil seeds were determined nonviable after the 4-day period (**Figure S1**, **Supporting Information)**. These results imply that 98% of the basil seeds used in this study can be germinated and accurate comparison of the differences in basil seed germination due to the experimental parameters can be investigated.

### Determination of the combined effect of Au NPs and microwave heating on basil seeds using optical microscopy and scanning electron microscopy

To investigate the effect of using 20 nm Au NPs and microwave heating on the basil seeds as compared to control samples (dry basil seeds and basil seeds in water without 20nm Au NPs/microwave heating), optical microscope images that show the top view of the whole basil seeds were taken before and after microwave heating or at room temperature without microwave heating (control experiment) (**Figure 2**). In this regard, basil seeds were exposed to the various conditions as described in the experimental section: dry basil seeds (no water, no Au NPs), basil seeds in water, in a solution of 20 nm Au NPs were kept at room temperature without microwave heating for 6 minutes and were exposed to continuous microwave heating at 2 W, 10 W and 20 W (maximum power of the microwave source at 8 GHz) for 6 minutes.

**Figure 2.**
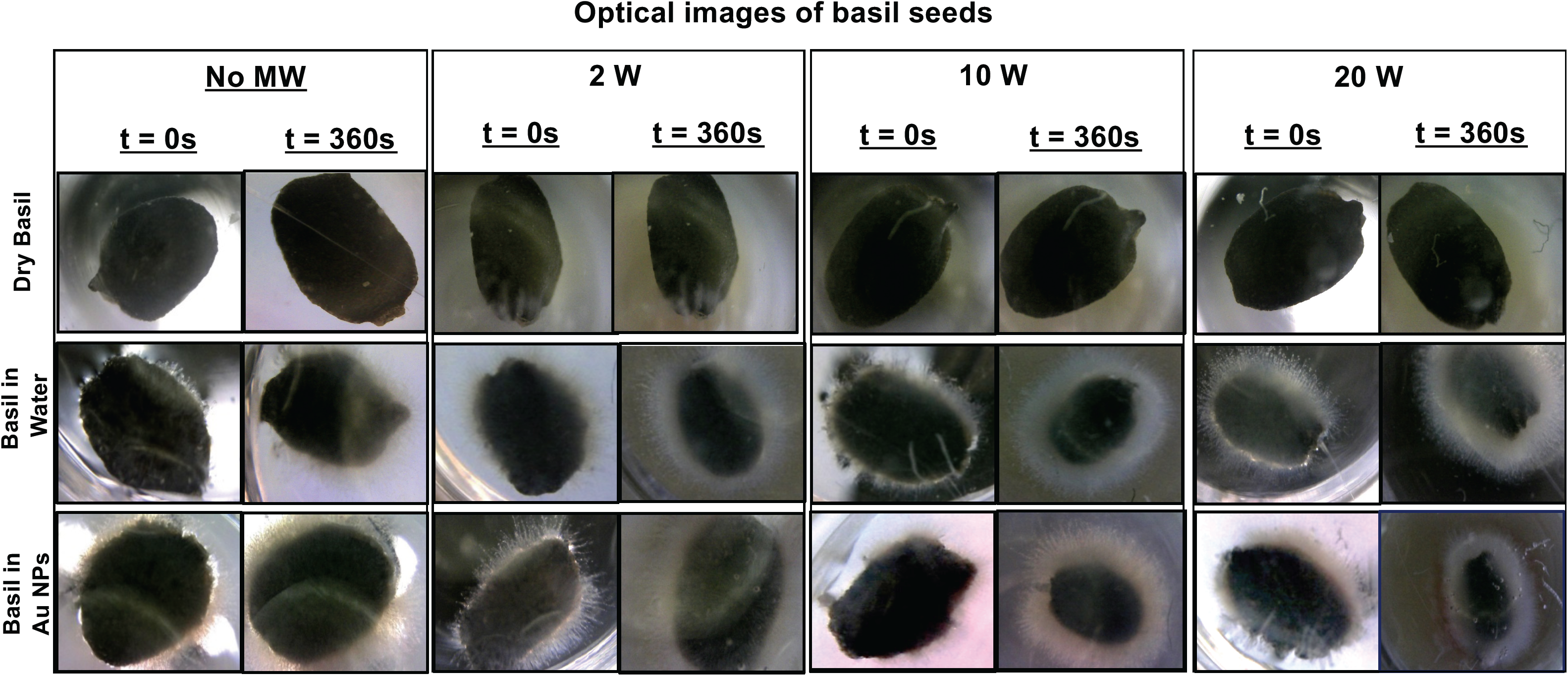
Bright-field images of basil seeds before and after application of microwave heating. Dry basil seeds, basil seeds in water (no Au NPs), or basil seeds in 20 nm Au NPs were microwaved at 2 W, 10 W, and 20 W of microwave power. The presence of Au NPs and microwave heating causes the seeds to develop longer gum within 6 minutes. Each basil seed is ca. 1.1 mm in length in the longest axis.

It is important to discuss the reasons for the selection of experimental conditions prior to the presentation of our observations. The choice for volume of water and 20 nm Au NPs solution (i.e., 50 μL sufficient to fully immerse up to 10 basil seeds) was due to the type of HTS well plates used (96 wells, flat-bottom for easy imaging from the bottom during microwave heating, maximum capacity of 100 μL). The choice for the type of metal nanoparticles (Au NPs, diameter = 20 nm) was due to the following factors: Au NPs are: 1) very stable in water, 2) increase enzyme activity, 3) act as “nano-bullets” in a microwave energy field, 4) easily observed individually under electron microscopy. The continuous microwave heating time of 6 minutes was chosen to minimize the changes to basil seeds in water at room temperature within 6 minutes observed using optical microscopy, therefore, the effect of 20 nm Au NPs and microwave heating on basil seeds can be clearly discerned while minimizing the time of exposure of basil seeds to microwave heating. Microwave heating of basil seeds longer than 6 minutes can potentially result in denaturation of enzymes in basil seeds due to increase in seed temperature >40°C and cause undesired structural damage to plant enzymes and germination process can be negatively affected.

#### Optical Microscopy

**Figure 2** shows the overall structural changes to the basil seeds observed using optical microscopy. In a control experiment, which was carried out to demonstrate that water is required to germinate basil seeds and that microwave heating of basil alone in the absence of water do not cause structural damage, no changes in the overall structure of the dry basil seeds was observed after 6 minutes at room temperature without microwave heating or with microwave heating. When kept in water and a solution of 20 nm Au NPs for 6 minutes at room temperature (without microwave heating), basil seeds developed a yellowish gum around the outer layer indicating the commencement of the germination process. Basil seeds kept in a solution of 20 nm Au NPs without microwave heating for 6 minutes developed longer whitish gum with red tint due to the presence of 20 nm Au NPs (solution of 20 nm Au NPs is red in color due their high scattering coefficients above 500 nm).

Microwave heating of basil seeds in water or in a solution of 20 nm Au NPs resulted in the growth of longer gum around the basil seeds. As the microwave power was increased from 2 W to 20 W, basil seed gum in water appeared to increase in length of and became dense and cloudier, which implies that microwave heating of basil seeds in water alone can increase the germination speed of basil seeds. Microwave heating of basil seeds in a solution of 20 nm Au NPs also resulted in the increase in length and change in the color of basil seed gum. These observations implied that Au NPs and microwave heating can be also used in increasing the speed of germination of basil seeds. Since microwave heating of dry basil seeds did not result in the growth of basil seed gum, the remainder of the experiments were carried out in water or in a solution of 20 nm Au NPs.

To investigate whether multiple basil seeds can be germinated at once using microwave heating, 10 basil seeds in water or in the presence of a solution of 20 nm Au NPs were placed in the same HTS wells (basil seeds were distributed at the bottom of the well without overlapping each other) and were exposed to continuous microwave heating at 2 W, 10 W and 20 W for 6 minutes. In addition, to monitor the structural changes in basil seeds during continuous microwave heating and to ascertain the extent of length of basil seed gum, optical images of basil seeds were captured from the bottom of the HTS wells every minute (**Figure S2**, **Supporting Information**). These observations reveal that the length of basil gum increases up to the 4^th^ minute of microwave heating and remains at the same length until the end of the experiment, that is, microwave heating of basil seed for 6 minutes is sufficient to initiate germination of basil seeds.

#### Scanning Electron Microscopy (SEM)

In addition to the optical microscopy images of basil seeds after microwave heating in the absence and presence of 20 nm Au NPs, extensive investigation of structural changes to the basil gum using SEM and elemental analysis was carried out (**Figure 3 and Figures S3-S12**, **Supporting Information)**, which provided an evidence for the overall structural changes the basil seeds. **Figures S3** shows the overall structural changes in the basil gum before (t = 0 min) and after each experimental condition (t = 6 minutes) observed by SEM (images acquired from four sides, top, left, right, bottom, for full view of the seeds). SEM images of dry basil seeds kept at room temperature without microwave heating show that seed surface is unchanged after 6 minutes. After 6 minutes of microwave heating of dry basil seeds, light-colored spots on the surface of seeds appeared due to microwave heating. Basil seeds placed in water without 20 nm Au NPs at room temperature without microwave heating for 6 minutes developed gum with a small gap on the top and bottom of the seeds. Microwave heating of basil seeds in water without 20 nm Au NPs for 6 minutes resulted in looser gum with large gaps around the seed surface, which indicates that microwave heating alone causes significant changes to basil seeds. Basil seeds placed in a solution of 20 nm Au NPs at room temperature without microwave heating resulted in observations similar to those observed for basil seeds kept at room temperature for 6 minutes. Microwave heating of basil seeds in a solution of 20 nm Au NPs for 6 minutes resulted in looser gum with large gaps around the seed surface. It is important to note that basil seeds were removed from water and the solution of 20 nm AuNPs to acquire the SEM images, therefore basil gum appears as wrapped around the seed rather than extending into the solution as seen in optical microscopy images.

**Figure 3.**
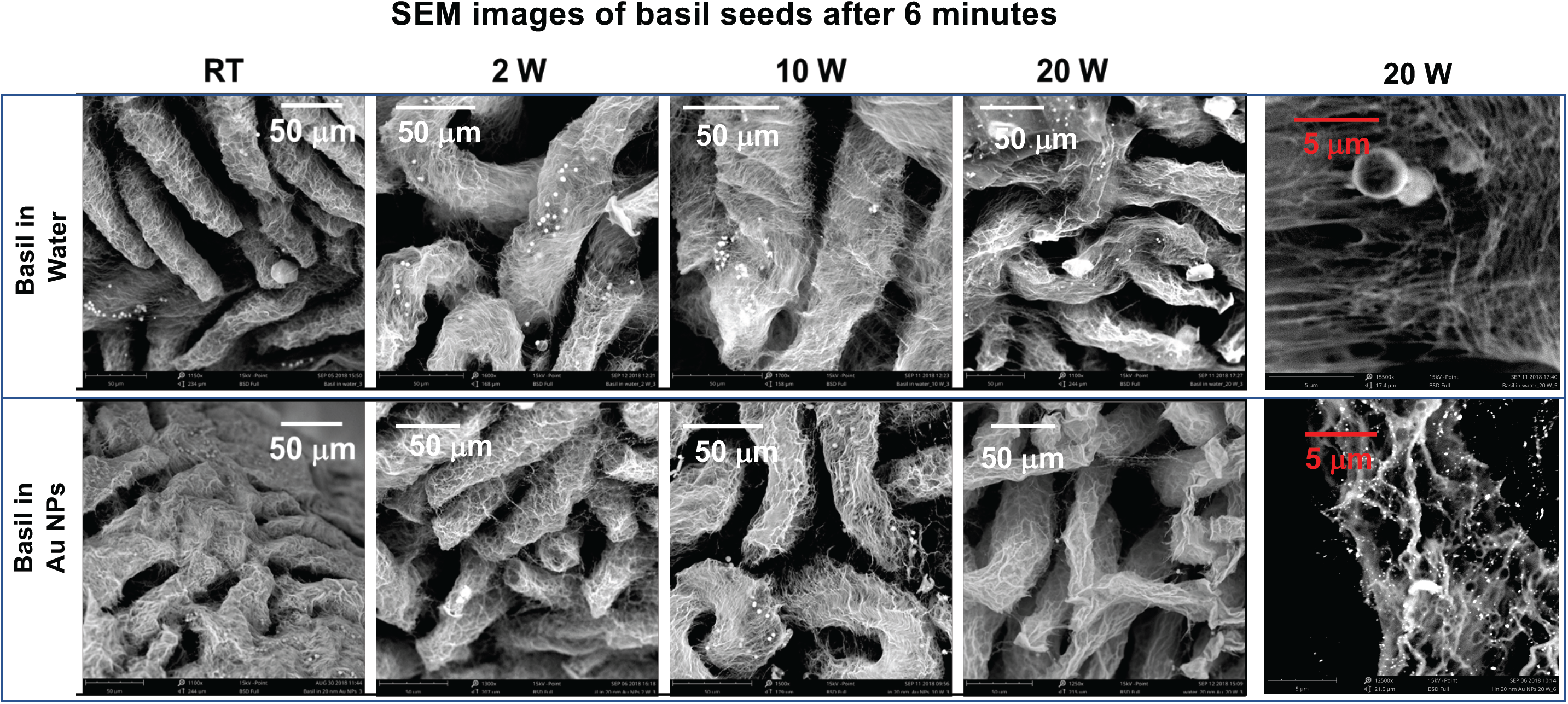
Scanning electron microscope images of basil seed gum after continuous microwave heating (2 W, 10 W, or 20 W) for 6 minutes. Control seeds were kept at room temperature for 6 minutes in either water or 20 nm Au NPs without microwave heating. Basil seed gum appears more spread out due to microwave heating and 20 nm Au NPs. Circular structures seen in basil in water is in 5 mm in size and part of the basil gum. Although Au NPs are significantly smaller than these circular structures, Au NPs are brighter due to their ability to scatter light more efficiently than the seed components. Scale bars = 50 mm and 5 mm (images on the right).

To gain further insight into the structural changes in basil seeds and discern the differences in the basil gum developed around the basil seeds in our experiments, SEM images of the basil gum after each experimental condition (except dry basil seeds, which lacked basil gum) were collected and compared. **Figure 3 (**also **Figure S5 and S9**, **Supporting Information)** show that basil gum appears more spread out in a solution of 20 nm Au NPs as compared to those kept in water. Microwave heating of basil seeds for 6 minutes results in a distribution of basil gum around the basil seeds depending on: 1) the level of microwave power and 2) the absence or presence of 20 nm Au NPs. Basil gum appears more spread out in basil seeds microwave heated as compared to basil gum kept in a solution of 20 nm AuNPs or water at room temperature without microwave heating. Similar observations are made as the microwave power is increased from 2 W to 20 W: basil gum appeared to spread out the most at 20 W, and the presence of 20 nm Au NPs increases the spreading of the basil gum. The observations discussed in the text so far imply the following: i) basil seeds develop basil gum faster in the presence of 20 nm Au NPs and microwave heating, and can germinate faster due to increased enzyme activity, or ii) basil gum is destroyed due to microwave heating, enzymes are denatured and basil seeds cannot germinate as compared to the control basil seeds (at room temperature, no Au NPs, no microwave heating).

In addition, circular structures *ca*. 5 μm in size that are part of the basil gum are seen around the basil seeds kept in water. Although 20 nm Au NPs are significantly smaller than these circular structures, 20 nm Au NPs are brighter due to their ability to scatter electrons more efficiently than the seed components. Individual 20 nm Au NPs are clearly visible and appeared to be distributed along the basil gum without aggregation as seen in SEM images, which can be attributed to the presence of enzymes (and other biological materials) in the basil seed and basil gum. Quantitative elemental analysis of the basil seeds after 6 minutes incubation in a solution of 20 nm Au NPs at room temperature or microwave heating provides evidence for the spatial distribution of 20 nm Au NPs around the basil gum (**Figure S5 – S12**, **Supporting Information)**. Since enzymes contain primary amine and thiol functional groups, Au NPs can chemisorb on to the enzymes through these functional groups. The effect of chemisorption of Au NPs on the esterase on the basil seeds is discussed in the **Supporting Information**.

Microwave heating of basil seeds in the presence of a solution containing 20 nm Au NPs at 2 W, 10 W and 20 W resulted in an increase of esterase activity as compared to esterase activity for basil seeds in water (**Figure S15**, **Supporting Information)**. These observations can be explained within the context of the combined effect of microwave heating on the basil gum and Au NPs-catalyzed esterase activity. (Deka *et al.*, 2012; Arsalan and Younus, 2018) As discussed in the previous paragraphs, microwave heating of basil seeds causes the basil gum to be looser as the microwave power is increased from 2 W to 20 W, and enzymatic activity is increased due to the availability of esterase around basil seed. In addition, microwave heating of basil seeds in the presence of 20 nm Au NPs changes the movement and distribution of 20 nm Au NPs around the basil gum, which in turn can affect the extent of interactions of 20 nm Au NPs with esterase in the basil gum. Our research group has previously shown that microwave heating of Au NPs in solution increases the diffusivity of Au NPs due to selective heating of water molecules in bulk and around the Au NPs.(Aslan and Geddes, 2007) Since the size of 20 nm Au NPs is *ca.* 1.88×10^8^ times smaller than the wavelength of microwaves at 8 GHz (3.75 cm) and the 20 nm Au NPs are negatively charged, 20 nm Au NPs are moved within microwave field without absorbing the microwave energy, while the temperature of water molecules around 20 nm Au NPs and in bulk is increased due to molecular friction, and subsequently, water is selectively heated. As a result of selective heating of water and coupling of negatively-charged 20 nm Au NPs with electromagnetic field, 20 nm Au NPs move about the basil seed faster and chemisorption events between 20 nm Au NPs and esterase are accelerated, which affects the distribution of Au NPs on the basil gum.

In this regard, the extent of temperature changes in bulk during microwave heating of basil seeds in water and 20 nm Au NPs solutions for 6 minutes and subsequent cooling period of 20 min taken to monitor enzymatic reactions was measured (**Figure S17**, **Supporting Information).** A maximum increase of *ca*. 11°C in temperature from room temperature (25°C) to 36°C was observed during microwave heating of basil seeds in water at 20 W. Microwave heating of basil seeds in a solution of 20 nm Au NPs at 20 W resulted in increase of temperature from room temperature to 34°C. Microwave heating of basil seeds in all solutions at 2 W and 10 W yielded high temperature in the range of 27°C-30°C, respectively. These observations imply that solutions are incrementally heated for 6 minutes below 37°C, where enzymes are not expected to denature due to microwave heating.

It is important to note that our experimental setup (**Figure S1**, **Supporting Information)** lacked the ability of simultaneous measurement of temperature (using a FLIR thermal camera) and enzyme activity (using a luminescence imaging instrument) during microwave heating of basil seeds. Therefore, enzymatic activity measurements were carried out immediately after 6 minutes of microwave heating of basil seeds for an additional 20 minutes. In addition, changes in temperature of solution after microwave heating was stopped for additional 20 minutes was measured to simulate the experimental conditions during enzymatic activity measurements. The temperature of all solutions returned to their initial values within 10 minutes. It is also important to note that after microwave heating for 6 minutes basil seeds were mixed with an equal volume of substrate solution kept at room temperature or directly transferred to a filter paper for hydroponic growth. The temperature of basil seeds returned to room temperature within 2 minutes after microwave heating and temperature does not affect enzyme activity after microwave heating was stopped. Therefore, we can conclude that esterase activity on basil seeds is affected mainly by microwave heating alone and microwave heating in the presence of 20 nm Au NPs for 6 minutes. Subsequently, microwave heating of basil seeds at 10 W was selected for hydroponic growth and soil growth experiments based on the following observations: i) temperature of the solution with basil seeds reaches a maximum value of 30°C when microwave heated at 10 W and ii) esterase activity can be modulated by using 20 nm Au NPs.

### Finite-Difference Time-Domain (FDTD) simulations and electrical conductivity measurements for basil seeds

In addition to experimental work to determine the effects of microwave heating of basil seeds, computational simulations were carried out to determine the percentage of microwave absorption by each component of the basil seed/water system and to visualize the electric field propagation through the system. **Figure S18 (Supporting Information)** shows the simulated electric field distribution around basil seeds with and without water (dry) exposed to monomode microwave point source operating at 8 GHz and 10 W. These simulations show that basil seeds in bulk water is predicted to absorb 9.1% of the homogenous electromagnetic energy. Conversely, dry basil seeds (no bulk water) are predicted to absorb a 0.7% extent of the electromagnetic energy due to the presence of water in basil seeds. The electric field propagation images also imply that the energy is absorbed by the basil seeds submerged in water, while the dry seeds transmit most of the incoming radiation.

To gain further insight to microwave heating of basil seeds through computational simulations of resistive losses of basil seeds (1, 3, 4, and 10 seeds) were carried out (**Figures 4 and S19-S20**, **Supporting Information)**. **Figure 4** shows the simulated resistive losses in terms of 12 cross-sections (in the xz-plane and y-direction) from four basil seeds in water exposed to monomode microwave point source operating at 8 GHz and 10 W using COMSOL software. Resistive losses are predicted to be similar for all four seeds and vary throughout all cross-sections of an individual basil seed. Since the microwave source is placed on top of the cavity, resistive losses from the top sections (slices 9-12) of the basil seeds are predicted to be the largest and the least resistive losses are predicted from the bottom sections (slices 1-4) of the basil seeds, while middle sections (slices 5-8) of the basil seeds show similar resistive losses. In addition, resistive losses from the top sections of the four basil seeds show the largest extent of losses occur from the sides of the basil seeds facing each other. In the middle and bottom sections of the basil seeds, resistive losses occur from the edges and opposite sides of basil seeds, respectively, which implies that the entire basil seed is heated. Therefore, basil gum is expected to develop from all sides of the basil seeds. Based on these computational simulations, resistive losses for a single seed, three seeds and ten seeds during microwave heating were also calculated (**Figures S19-S20**, **Supporting Information)**, which predict similar patterns for resistive losses as described for four basil seeds.

**Figure 4.**
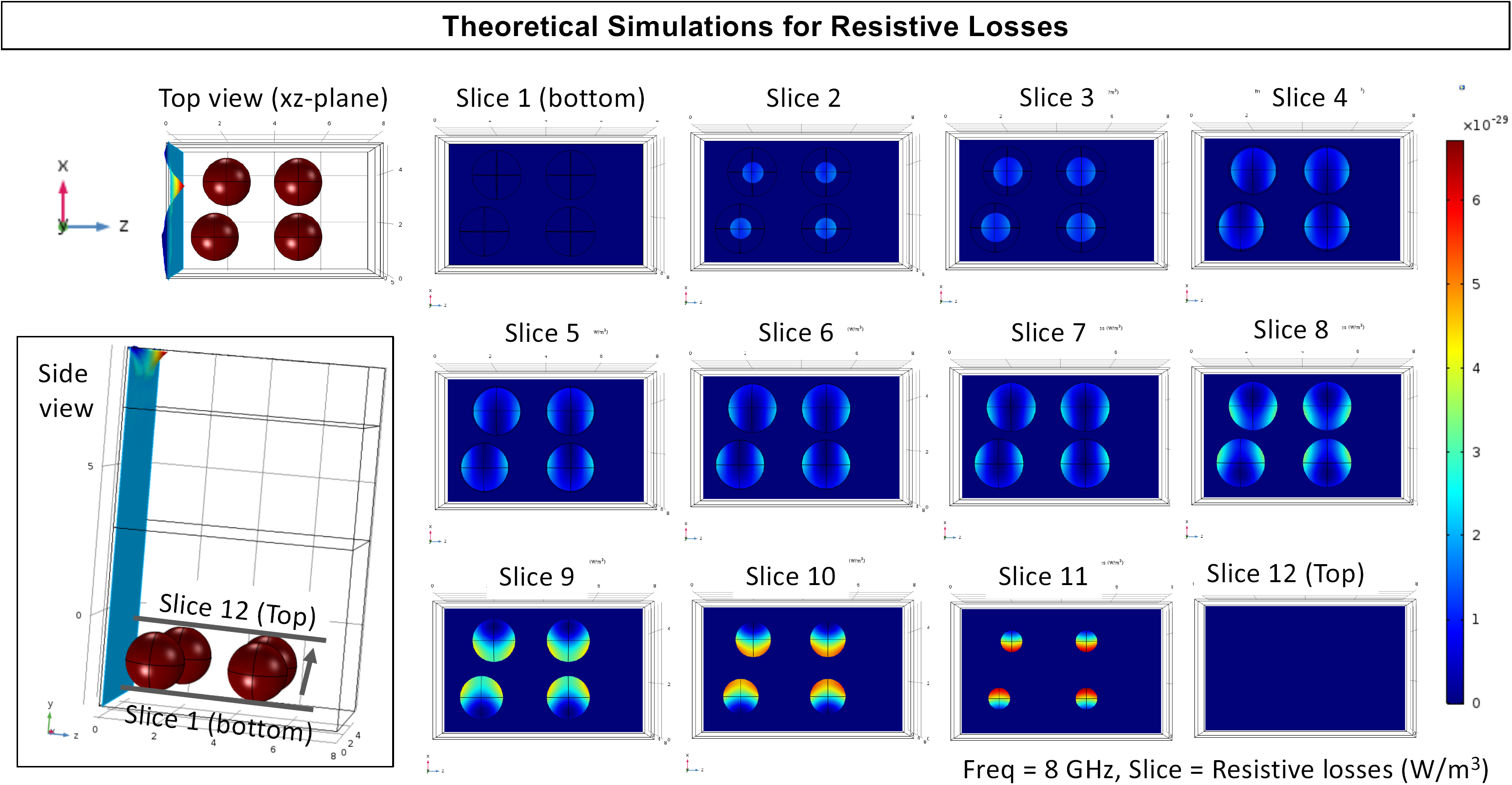
FDTD simulation of resistive losses from basil seeds fully immersed in water exposed to monomode microwave point source operating at 8 GHz and 10 W using COMSOL software. Top and side view of the microwave cavity and seeds are shown on the left. Resistive losses from the seeds are shown as slices starting from the layer bottom of the seeds up to the layer corresponding to the plane on top of the seeds.

In addition to theoretical calculations, to further our understanding of the effect of microwave heating on basil seeds, electrical conductivity of basil seeds immersed in water and in a solution of 20 nm Au NPs for 6 minutes was measured to be: 0.140 S/(cm.g) for basil seeds in water and 0.156 S/(cm.g) for basil seeds in 20 nm Au NPs. Since the resistivity of dry basil is extremely high, no electrical conductivity values were measured. These experimental measurements show that basil seeds in water and in a solution of 20 nm Au NPs have finite electrical conductivity, and electrical component of microwave energy can couple to the basil seeds and be converted into heat through resistive losses in the basil seeds. Since the bulk temperature of solutions containing basil seeds only reach a maximum value of 30°C when microwave heated at 10 W, we can conclude that the development of basil gum is accelerated due to bulk heating and resistive losses in the seeds while enzyme activity is maintained. In the absence of microwave heating, development of basil gum in water is slower at room temperature, as shown in optical images in this study.

### Effect of gold nanoparticles and microwave heating on hydroponic growth of basil seeds

Based on our ability to control the development of basil gum using 20 nm Au NPs and microwave heating as described above, we investigated the effect of 20 nm Au NPs and microwave heating on hydroponic growth of basil seeds to answer the question of “*Can the use of 20 nm Au NPs and microwave heating result in modulation of hydroponic growth of basil seeds*?”. In this regard, basil seeds immersed in water or in a solution 20 nm Au NPs were microwave heated at 10 W for 6 minutes and placed in a hydroponic growth setup as described in the experimental section of this study. In addition, control experiments, where basil seeds were kept at room temperature without microwave heating in water and a solution of 20 nm Au NPs, were also carried out to discern the differences in using microwave heating versus basil seed growth at room temperature without microwave heating.

To establish a reference point for the hydroponic growth experiments with microwave heating treated basil seeds, we first discuss hydroponic growth from basil seeds kept at room temperature without microwave heating. **Figure 5** (see also **Figure S21**, **Supporting Information)**, shows that basil seeds kept in water at room temperature for 6 minutes (i.e. control sample) developed long stems at 72 hours and grow to full reference length with leaves at 128 hours. At 72 hours, basil seeds kept in a solution of 20 nm Au NPs at room temperature grew shorter stem as compared to control sample, which implies that the development of basil plant is delayed in the presence of 20 nm Au NPs. At 72 hours, basil seeds exposed to microwave heating in the presence of 20 nm Au NPs (i.e., sample labeled as MAMAG) developed longer stem and leaves as compared to the other samples, which implies that the development of basil plant is accelerated with the combined use of 20 nm Au NPs and microwave heating. At 128 hours, basil plants grown from basil seeds treated with the MAMAG technique (i.e., combined used of Au NPs and microwave heating) displayed the largest leaves and longest stem, while the growth of basil plants grown from basil seeds kept in the solution of 20 nm Au NPs remained delayed and the control sample reached its peak value in stem length and leaf size. At 224 hours, two of the three basil plants grown from basil seeds treated with the MAMAG technique were still viable without any sign of deterioration of leaves or the plant stem. At 224 hours, control sample showed significant deterioration of leaves and the plant stem and one of the three basil plants grown from basil seeds kept in the solution of 20 nm Au NPs remained viable. At the end of our observations of basil plant growth at 328 hours, all basil plants deteriorated completely. These observations clearly demonstrated that the hydroponic growth of basil plants can be delayed with the use of 20 nm Au NPs without microwave heating and accelerated and extended with the combined use of 20 nm Au NPs and microwave heating.

**Figure 5.**
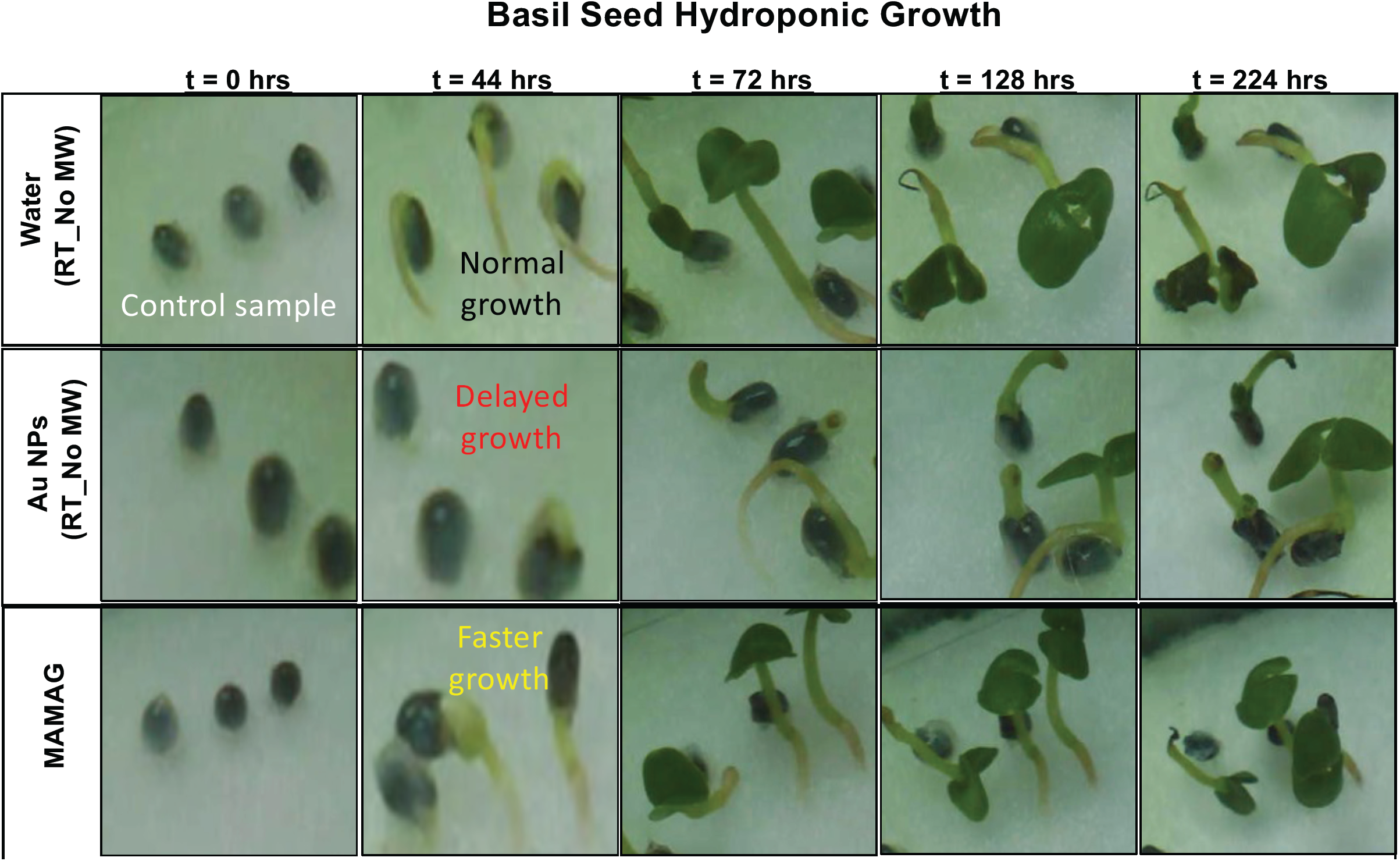
Basil seeds in hydroponic growth. Three basil seeds were kept in water without Au NPs at room temperature (No MW/RT), or kept in a solution of 20 nm Au NPs at room temperature, or microwave heated (MW) in in a solution of 20 nm Au NPs for 6 min at 10 W and placed into a modified growth system. Basil seeds in water at room temperature (no microwave heating) are used as a control sample to monitor the normal hydroponic growth.

### Effect of gold nanoparticles and microwave heating on growth of basil seeds in soil

In the final step of this study, we investigated the effect of 20 nm Au NPs alone and combined with microwave heating on the growth of basil seeds to find out whether the growth of basil seeds in soil can be delayed or accelerated on demand. In this regard, after incubation for 6 minutes at room temperature in water or 20 nm Au NPs or in 20 nm Au NPs with microwave heating (at 2 W, 10 W, and 20 W), basil seeds were placed in seedling starter trays to monitor growth in soil for up to 73 days in identical conditions. After 5 days of planting, basil seeds kept in water at room temperature for 6 minutes (i.e., no microwave heating) developed two-leaves, which was used as a reference point (control sample) for the growth of basil seeds in the presence of 20 nm Au NPs and microwave heating (**Figure 6**). Basil seeds kept in in 20 nm Au NPs at room temperature without microwave heating developed smaller leaves and the height of the basil plants were shorter than those observed for the control sample, which implied that the growth of basil plants from their seeds treated with 20 nm Au NPs at room temperature was delayed in soil. Microwave heating of basil seeds (at 2 W, 10 W, and 20 W) in the presence of 20 nm Au NPs for 6 minutes resulted in taller basil plants with wider leaves after 5 days of planting in soil, which implied that the growth of basil plants from their seeds microwave heated in the presence of Au NPs was accelerated in soil. Furthermore, after 73 days planting of basil seeds, significant differences between basil plants were observed: while basil plants grown from basil seeds kept in water at room temperature (i.e., no microwave heating) for 6 minutes wilted, basil plants grown from basil seeds kept in a solution of 20 nm Au NPs at room temperature and basil seeds microwave heated in a solution of 20 nm Au NPs were still thriving 73 days after planting in soil (**Figure 6**). It is important to note all basil plants were exposed to identical experimental conditions. Moreover, the height of the basil plants grown from basil seeds kept in a solution of 20 nm Au NPs at room temperature were taller than any basil plant grown under other conditions.

**Figure 6.**
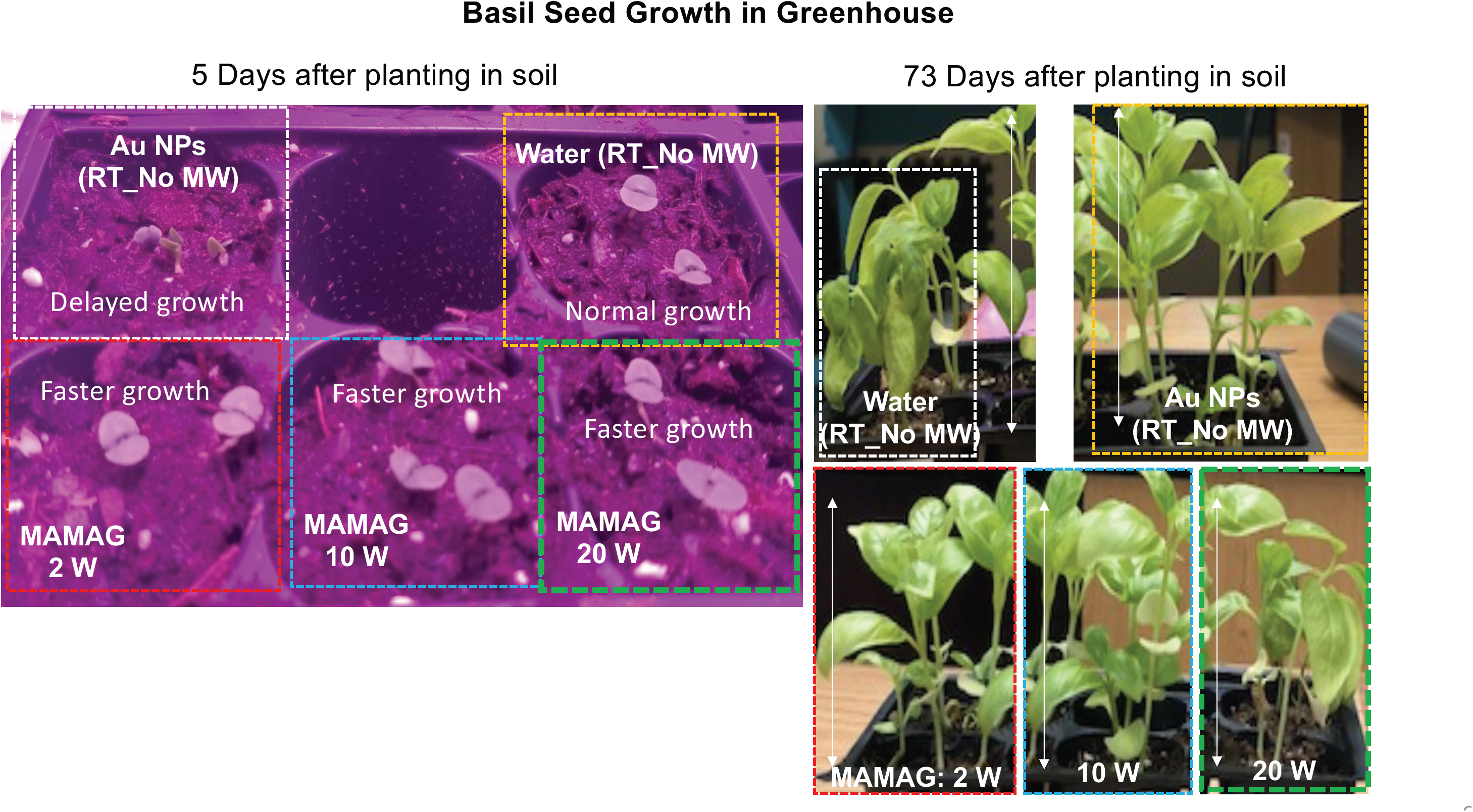
Basil seed growth in soil in a greenhouse. Basil seeds were microwaved continuously for 6 minutes at 2 W, 10 W, and 20 W. Basil seeds in water only (no microwave heating) and basil seeds in 20 nm Au NPs only (no microwave) were used as control basil seeds. All pictures were taken together with the same camera. Vertical white line shows the length of basil plants = 7 cm.

Since microwave heating of basil seeds in water and in 20 nm Au NPs results in an increase in temperature up to 36°C, we investigated the effect of incubation of basil seeds in pre-heated water without microwave heating to compare these results with the results obtained using basil seeds treated with microwave heating. **Figures S22** (**Supporting Information)** show that the development of basil gum, enzymatic activity and soil growth in 17 days were similar to the results observed for basil seeds kept at room temperature without microwave heating, and the MAMAG technique, based on combined use of microwave heating and 20 nm Au NPs, for the pre-treatment of basil seeds offer a superior alternative to simply soaking basil seeds in pre-heated water. These observations imply that the growth of basil seeds in soil can be delayed or accelerated on demand with 20 nm Au NPs and microwave heating.

## CONCLUSIONS

In this work, we demonstrated that a solution of 20 nm Au NPs and microwave heating from a monomode microwave source can be used as new approach to modulate the germination of basil seeds and subsequent hydroponic growth and soil growth of basil plants. In this regard, single basil seeds or 10 basil seeds were placed in a 50 μL solution of 20 nm Au NPs in a HTS well plates and were exposed to 6 minutes of continuous microwave heating using a solid-state 8 GHz microwave generator operating at 2 W, 10 W and 20 W microwave power. In control experiments, either 20 nm Au NPs or microwave heating was omitted to determine the effect of each of 20 nm Au NPs alone (at room temperature without microwave heating) or microwave heating alone (in water without 20 nm Au NPs). We have made the following conclusions from our experimental observations and computational simulations:

1. Optical microscopy was used to assess the macro-scale changes in basil seeds and showed that
  a. dry basil seeds did not germinate with or without microwave heating, and
  b. basil in water or in a solution of 20 nm Au NPs in the presence of microwave heating resulted in longer basil gum as compared to basil seeds kept in room temperature without microwave heating.
2. SEM was used to assess the micro-scale changes in basil seed gum and showed that
  a. basil seed gum is more spread out due to microwave heating and in the presence of 20 nm Au NPs, and
  b. 20 nm Au NPs are distributed throughout the surface of the basil seeds without significant aggregation.
3. Luminescence kinetic analysis of enzymatic activity was used to assess the effect of 20 nm Au NPs and microwave heating on esterase, which is a key enzyme for germination of basil seeds. Esterase activity was monitored for an additional 20 minutes after the initial 6 minutes incubation of basil seeds,
  a. esterase activity in basil seeds in water and in a solution of 20 nm Au NPs without microwave heating was similar, and
  b. esterase activity in basil seeds in 20 nm Au NPs was up to 2-fold higher than esterase activity in water with microwave heating.
4. Maximum actual average temperature of water and 20 nm Au NPs containing basil seeds during microwave heating at 20 W was measured to be 36°C and 34°C, respectively. Enzymes in basil seeds are not expected to denature due to microwave heating of basil seeds in water and in a solution of 20 nm Au NPs.
5. FDTD simulations predict that
  a. basil seeds in water absorbs 9.1%, transmits 85.4%, and reflects 5.5% of the microwave radiation.
  b. Resistive losses from the top sections of the basil seeds show the largest extent of losses occur from the sides of the basil seeds facing each other. Resistive losses occur from the edges and opposite sides of basil seeds in the middle and bottom sections of the basil seeds, respectively.
  c. The entire basil seed is heated during microwave heating, and therefore basil gum is expected to develop from all sides of the basil seeds after microwave heating.
6. Hydroponic growth of basil plants can be
  a. delayed with the use of 20 nm Au NPs at room temperature without microwave heating, or
  b. accelerated and life of the basil plant can be extended with the combined use of 20 nm Au NPs and microwave heating.
7. Growth of basil seeds in soil can be
  a. delayed by pre-treatment of basil seeds in a solution of 20 nm Au NPs, or
  b. accelerated after pre-treatment of basil seeds with microwave heating and a solution of 20 nm Au NPs.

## ACKNOWLEDGMENTS

Authors greatly appreciate partial financial support from Morgan State University Innovation Works I-GAP Program.

## SUPPORTING INFORMATION

Additional SEM images, luminescence microscopy images of basil seeds, detailed hydroponic growth results, real-time temperature measurements of the well with a single basil seed at room temperature and FDTD simulation of resistive losses are presented.

---

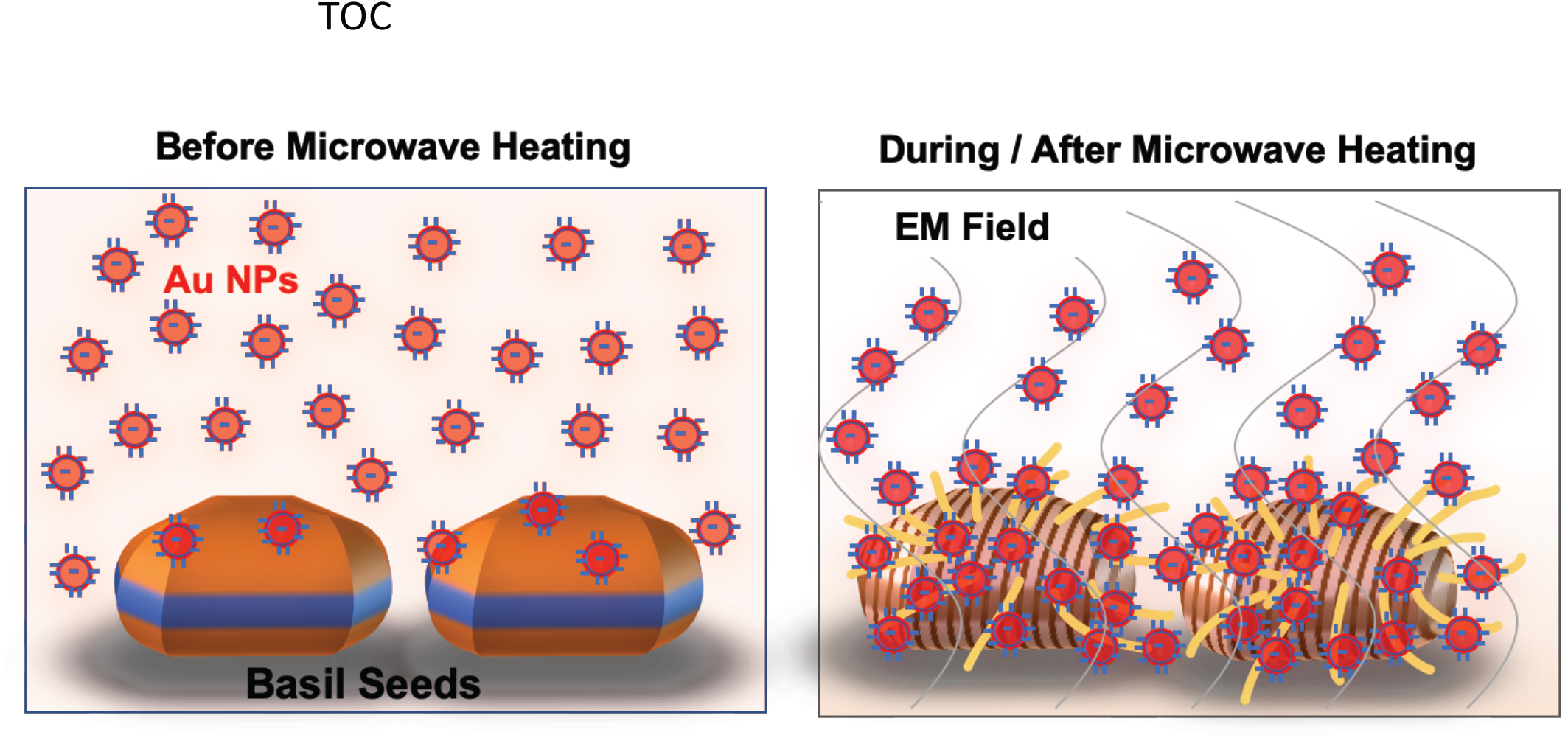

